# Heartfelt gaze: Cardiac afferent signals and vagal tone affect gaze perception

**DOI:** 10.1101/2025.01.15.632948

**Authors:** Yaojie Lin, Tomoko Isomura

## Abstract

Perceiving others’ gaze direction is an essential aspect of social interactions. The cone of direct gaze (CoDG) refers to the range within which an observer perceives gaze as looking directly at them. Previous research has demonstrated that self-relevant exteroceptive cues can widen CoDG. However, the effect of self-relevant interoceptive information on the CoDG remains unknown. In the present study, we used a modified gaze discrimination task to synchronize face stimuli with various gaze directions to specific phases of the cardiac cycle. The results showed that participants with higher heart rate variability (HRV) exhibited a wider CoDG during cardiac systole (when cardiac signals are maximally represented in the brain), but not during cardiac diastole (when these signals are quiescent). Moreover, this effect was independent of individual differences in anxiety levels and autistic traits. We interpret these findings as evidence that individuals with greater cardiac vagal control are more sensitive to cardiac afferent signals during systole, which in turn leads to a stronger self-directed perception of others’ gaze under transient and ambiguous gaze perception conditions. Our findings highlight the self-referential role of cardiac interoceptive signals in gaze perception, expanding our knowledge of interoceptive influences on social judgment.

## 1. Introduction

Perceiving the gaze of others is an important social skill in our daily life. It underlies the promotion of social interactions with others and plays a significant role in the development of social cognition (Baron-Cohen, 1995; Kendon 1967; Kleinke 1986). Gaze direction in others often signals interest and approach orientation but can also indicate potential threats in the environment (Baron-Cohen, 1995; Hadjikhani et al., 2008; Lobmaier et al., 2021). The ability to rapidly detect gaze direction may reflect a highly adaptive mechanism that is likely to be critical for survival (Fox et al., 2007). For instance, the ability to distinguish between direct and averted gaze and preference for direct eye contact is evident from birth (Farroni et al., 2002). Moreover, studies have demonstrated that the perception of others’ gaze direction facilitates early visual processing, with neural responses and reflexive orienting being triggered in the 100 ms range following the onset of the gaze stimulus (Friesen & Kingstone, 1998; Schuller & Rossion, 2004; Schuller & Rossion, 2005).

When detecting the gaze from others, however, we do not interpret it as a precise “ray” but rather as an overly sensitive perception of a vague “range”, where a considerable span of averted gaze can also be perceived as directed at the observer, commonly known as the cone of direct gaze (CoDG; Gamer & Hecht, 2007). A wider CoDG suggests that the observer makes more liberal judgments (Gianotti et al., 2018). Although the width of an individual’s CoDG has been reported as a relatively stable trait (Lobmaier et al., 2021), there is significant variability between individuals in the width of the CoDG. People with higher anxiety levels tend to exhibit a wider CoDG compared to those with lower anxiety levels (Chen et al., 2017; Hu et al., 2017; Schulze et al., 2013). On the other hand, individuals with higher autistic traits tend to have a narrower CoDG (Matsuyoshi et al., 2014; but see Williams et al., 2023).

Research on gaze effects have shown that direct gaze trigger self-referential processing (Hietanen & Hietanen, 2017; for a review, see Conty et al., 2016) and enhances self-awareness (Argyle, 1975; Baltazar et al., 2014; Hazem et al., 2017; Hietanen et al., 2008; Isomura & Watanabe, 2020; Pönkänen et al., 2011). On the other hand, current research suggest that self-relevant information and self-referential processing can conversely widen the range of feeling being looked at directly by others. Stoyanova and colleagues (2010) demonstrated that the CoDG widened when participants heard their own names during the presentation of neutral face stimuli, compared to hearing someone else’s name. Similarly, Chen and Hietanen (2024) showed that participants perceived a broader range of gaze deviations as direct eye when the speech was directed to them as opposed to when it was directed to others. Moreover, it has also been proposed that increased self-referential processing in psychopathology may contribute to the widening of gaze cone (Hietanen et al., 2022; Wastler & Lenzenweger, 2018). Furthermore, facial emotions of others have also been shown to modulate the CoDG by induing self-referential processing. For instance, angry faces are thought to be looking at the participant over a wider range of gaze directions, which may reflect an adaptive response to interpersonal threat, driven by a self-referential bias that prioritizes recognizing potential danger in ambiguous situations (Ewbank et al., 2009; Harbort et al., 2013; Rhodes et al., 2012), while some studies show that happy faces were more likely to be judged as looking at the observer than other faces, which has been interpreted by the self-referential positivity bias, suggesting a general inclination to associate other people’s happiness with the self (Lobmaier et al., 2008; Lobmaier & Perrett, 2011). Collectively, these findings provide evidence that self-referential processes triggered by exteroceptive information (visual and auditory) can expand the subjective range of perceived direct gaze.

The construction and maintenance of our understanding of the world is driven by the coordinated integration of exteroceptive and interoceptive processes. Interoception refers to the process of detecting, interpreting, and integrating the internal states of the body across conscious and unconscious levels (Khalsa et al., 2018). The majority of studies to date have focused on cardiac interoception due to its discreteness, ease of measurement, and quantifiable performance accuracy (Garfinkel et al., 2016). Cardiac activity can be divided into two main phases: diastole (the ventricles expand and fill with blood) and systole (the ventricles contract and eject blood into the arteries). Baroreceptors are the pressure and stretch sensors in the aortic and carotid arteries, transmitting information about the heartbeats to the brain via afferent pathways in systole across the cardiac cycle. A large body of studies has shown that cardiac afferent signals exert pervasive influences on cognitive processes (for reviews, see Critchley & Garfinkel, 2015; Skora et al., 2022), such as perception (Edwards et al., 2002; Galvez-Pol et al., 2022; Saltafossi et al., 2023; Wilkinson et al., 2013), emotion (Gray et al., 2012; Garfinkel et al., 2014; Garfinkel & Critchley, 2016; Pfeifer et al., 2017), memory (Fiacconi et al., 2016; Garfinkel et al., 2013), decision-making (Kimura et al., 2023; von Mohr et al., 2023).

Recent empirical evidence implies the possibility that baroreceptor activation during systole set the physiological context of self-awareness and elicit self-referential processing in social judgement (Ambrosini et al., 2019; Honda & Nakao, 2022; von Mohr et al., 2021). In a self-face recognition task, Ambrosini et al. (2019) showed that presenting a single image in synchrony with the participant’s systole facilitated the self-face processing relative to diastole, indicating that baroafferent information speeds up self-recognition. Furthermore, studies have also indicated that systole is associated with egocentric bias and self-prioritization effects, particularly in individuals with heightened interoceptive accuracy (i.e., the ability to accurately perceive internal bodily signals). For example, von Mohr et al. (2021) reported that the emotional egocentricity bias (EEB) is stronger at the systolic versus the diastolic condition, revealing a tendency to enhance one’s own emotional experiences as a reference when judging other’s emotional state during baroreceptor activation. Honda and Nakao (2022) conducted a shape-label matching paradigm and found a reduced discrimination of self-relevant stimuli when newly self-associated geometric shapes were presented during systole, suggesting a preferential processing of internal self- relevant information over external self-relevant information during systole. Baroreceptor activation during systolic phase conveys information about internal bodily states, represents a crucial period of body-brain interactions (Critchley & Harrison, 2013) and the neural responses to cardiac inputs (i.e., heartbeat-evoked activity) has been associated with self-consciousness and self-referential processing (Babo-Rebelo et al., 2016; Park et al., 2016; Park et al., 2018; Park & Blanke, 2019).

So far studies have only focused on the contribution of self-related exteroceptive information to gaze perception, however, evidence regarding the self-related interoceptive mechanism underlying this effect remains scarce. The present study aimed to investigate the contribution of cardiac afferent signals and cardiac vagal tone to the perception of gaze from others. We performed a modified gaze discrimination task, programmed to deliver face stimuli with different gaze directions synchronized to either the systolic or diastolic phases of the participant’s heartbeat, calculating the width of the CoDG in different cardiac phases separately. We hypothesize that presenting stimuli during the systolic phase may widen the CoDG due to its physiological context of self-awareness compared to the diastole. Heart rate variability (HRV) is an index of cardiac vagal tone (Laborde et al., 2017), generated by heart-brain interactions and reflect the strength of parasympathetic influence on cardioregulation (Shaffer & Ginsberg, 2017; Thayer et al., 2012). Recent studies on cardiac effects have suggested that individual differences in the effects of cardiac cycle on cognition that may be attributed to individual differences in the cardiac vagal tone (e.g., Izaki et al., 2024; Rae et al., 2018), since vagally mediated HRV is thought to be closely linked to parasympathetic activation via baroreceptor firing (Azevedo et al., 2018). Hence, we estimated the resting-state HRV to investigate the influence of the cardiac vagal tone on cardiac cycle modulation of gaze perception. In addition, we measured the participants’ anxiety levels and autistic traits considering their potential effects on CoDG.

## 2. Methods

### 2.1. Participants

Forty students from Nagoya university (25 females; mean age = 21.48 years, SD = 2.65 years) participated in the study. The required sample size was calculated a priori using G*Power 3.1 (Faul et al., 2007), based on an α error of 0.05, a power of 0.8, and an effect size of Cohen’s d = 0.5 (medium-sized effect; Cohen, 1998), indicating that at least 34 participants would be necessary to detect significant within-subjects effects. All participants had normal or corrected to normal vision. Autistic traits and anxiety levels were assessed using the Autism-Spectrum Quotient (AQ) (Baron-Cohen et al., 2001; Japanese version by Wakabayashi et al., 2004) and the State-Trait Anxiety Inventory (STAI) (Spielberger et al., 1970; Japanese version by Shimizu & Imae, 1981). Three participants’ data were excluded from the analysis due to irregular cardiac rhythms, one was excluded for not meeting the experimental requirement of a quiet environment, and another was excluded due to exceptionally low response accuracy for direct (61%) and 9° averted gaze (42%). Thus, a total of 35 participants (22 females; mean age = 21.61 years, SD = 2.73 years) were included in the final analyses. This study was approved by the Ethics Committee of Nagoya University (No. NUPSY-240729-L-01). All participants provided written informed consent.

### 2.2. Stimuli

The face stimuli consisted of four Japanese (two males and two females) portrayed neutral expressions were selected from the Kokoro Research Center (KRC) facial expression database (Ueda et al., 2019). All necessary permissions to use these images in the current study and for preprint publication on bioRxiv have been obtained. Using Adobe Photoshop, we converted the photographs to grayscale, manipulated the gaze by shifting the position of each eye’s iris in 1-pixel increments per image (equivalent to a shift of 0.03 cm), and masked non-facial attributes (hair, ears, background), leaving only the central face area visible. The facial images subtended a visual angle of 11.73° by 8.86° on screen at a viewing distance of 60 cm. We used a total of seven gaze deviations for each face stimulus: true direct gaze plus shifts of 3, 6, and 9 pixels for left and right gaze (see Fig. 1). These deviations were chosen based on previous studies of the cone of direct gaze (e.g., Hu et al., 2017; Lebert et al., 2021). A total of 28 facial images were used for the experiment (4 faces with 7 gaze deviations).

**Fig. 1.**
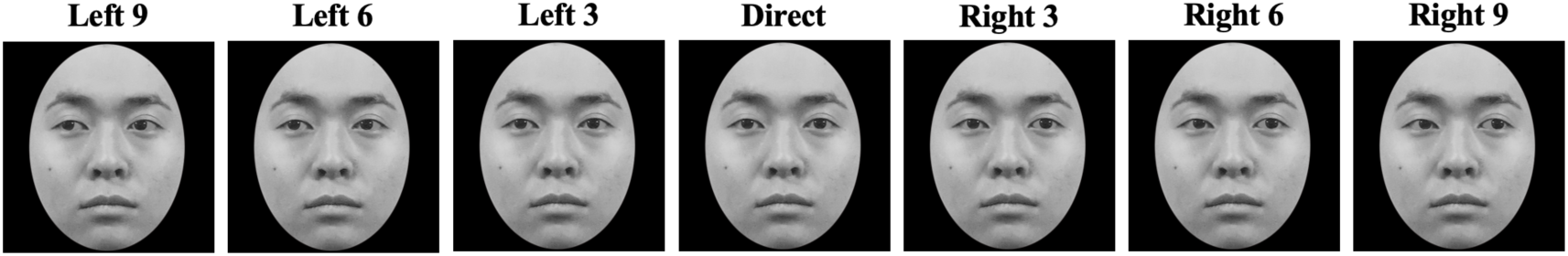
Examples of the face stimuli. A male frontal face with a neutral expression was shown at seven gaze deviations: 9 pixels left, 6 pixels left, 3 pixels left, direct gaze, 3 pixels right, 6 pixels right, and 9 pixels right. The photographs used in this figure were obtained from the Kokoro Research Center (KRC) facial expression database (Ueda et al., 2019) and permission for use has been obtained from the copyright holders.

### 2.3. Materials and Apparatus

The experiment was conducted using the MATLAB 2021b (Mathworks, Natick, MA, USA) and the Psychtoolbox-3 (Kleiner et al., 2007). The stimuli and instructions were centrally presented on a 23.8- inch PC monitor with a refresh rate of 60 Hz and a resolution of 1920 × 1080 pixels.

The Electrocardiography (ECG) data was recorded throughout the experiment using a BIOPAC MP160 system with a BIOPAC ECG100C amplifier (BIOPAC Systems, Goleta, CA) and three Ag/AgCl electrodes. The ECG signal was recorded using a 2000 Hz sampling rate and a 1 to 35 Hz band-pass filter. We used the DTU100 Digital Trigger Unit for the real-time detection of R waves in the ECG signal to synchronize face stimulus presentation to different phases of the cardiac cycle.

### 2.4. Procedure

Upon arrival at the laboratory, participants provided written informed consent and completed questionnaires consisting of the Japanese version of the Autism-Spectrum Quotient (AQ; Wakabayashi et al., 2004), the state scale of the State-Trait Anxiety Inventory (STAI-s) and the trait scale of the State-Trait Anxiety Inventory (STAI-t) (Shimizu & Imae, 1981). ECG electrodes were then attached underneath the left and right collarbone and one to the right side of the waist. Participants sat approximately 60 cm from the screen with their chin on a chin rest. The gaze discrimination task was conducted in a dimly lit and quiet room.

In a within-subjects design, we synchronized the onset of presentation of the face stimuli to either the systolic phase (defined as 200 ms after the R peak) or to the diastolic phase (500 ms after the R peak) of the cardiac cycle (Azevedo et al., 2023; Herman & Tsakiris, 2021; von Mohr et al., 2023) for a duration of 100 ms, ensuring that stimuli occurred at specific points in the cardiac cycle (Garfinkel et al., 2014). Participants received instructions on screen and performed a practice session consisting of 21 trials prior to the main experimental task. The main task consisted of 224 trials (112 trials for both the systolic and diastolic conditions) divided into four blocks of 56 trials. Within each block, participants were randomly assigned to either a systolic or a diastolic trial, with the constraint that each face identity and gaze angle was distributed across the blocks equally. Between each experimental block, participants were given a 30-second break.

The trial started with a fixation cross, with a total duration ranging between 1000 and 3000 ms, determined by the detection of the first R wave after a fixed 1000 ms period. The face stimulus was then presented coincide with cardiac systole or with cardiac diastole for 100 ms of the participant’s heartbeat. Following the stimulus, a blank screen was shown for 200 ms, after which the question ‘Where was the face looking?’ appeared in the center of the screen for up to 2000 ms. Participants were instructed to indicate as quickly and as accurately as possible where the presented face was looking during the response window. Responses were entered using the ‘J’, ‘K’, and ‘L’ keys of a standard QWERTY keyboard (for left-handed participants, the ‘A’, ‘S’ and ‘D’ keys were used instead) to indicate leftward, direct and rightward gaze. If no response was made within the 2000 ms time frame, the next trial began after a 500 ms inter trial interval (ITI) (see Fig. 2).

**Fig. 2.**
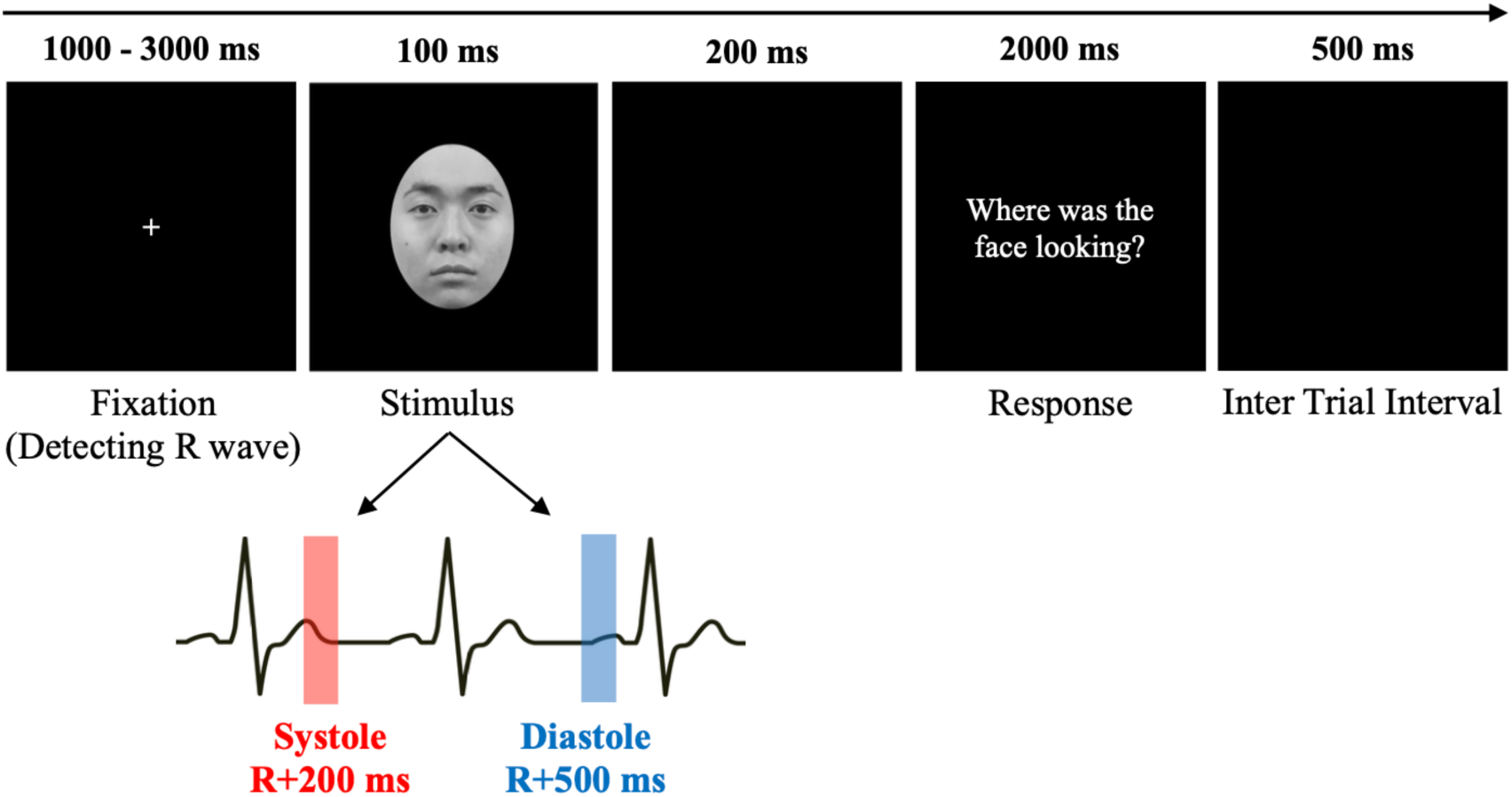
Illustration of the experimental trial in the gaze discrimination task. The face stimulus was either synchronized with cardiac systole (200 ms after the R peak) or cardiac diastole (500 ms after the R peak).

After the gaze discrimination task, participants were seated comfortably in chair and asked to rest for 5 min. During this period the ECG was recorded in order to estimate the participants’ resting HRV.

### 2.5. Statistical analyses

Statistical analyses were performed using Matlab 2021b (Mathworks, Natick, MA, USA) and JASP version 0.19.0 (JASP Team, 2024) (https://jasp-stats.org/). Movement or irregularities in heart rate during the experiment may introduce artifacts, leading to inaccurate R-wave detection and compromising the precise presentation of stimuli at specific points in the cardiac cycle. Therefore, prior to analyzing the behavioral data, both visual inspection and in-house Matlab scripts were used to detect and exclude trial data affected by such artifacts, as well as trials with no response (missing response data).

The CoDG were measured using methods similar to those in previous studies (Ewbank et al., 2009; Hu et al., 2017; Stoyanova et al., 2010). For each participant’s data, logistic functions were fitted to the proportions of “left” and “right” responses and the function representing “direct” responses was derived by subtracting the sum of “left” and “right” responses from one. Based on these data, the crossover points between the “direct” and “left” curves and between the “direct” and “right” curves were identified. The CoDG was calculated as the sum of the absolute values of these two crossover points (in pixels), representing the range of gaze deviations perceived as direct by each participant. The process was conducted separately for each cardiac condition (systole vs. diastole).

Heart rate variability (HRV) indices were computed based on the 5-minute ECG recording of resting period. We computed the root mean square of successive differences (RMSSD) from the R–R intervals as a time-domain index of HRV. The low-frequency HRV (LF-HRV; 0.04 – 0.15 Hz) and high-frequency HRV (HF-HRV; 0.15 – 0.4 Hz) components were extracted from the power spectral density (PSD) of HRV using the Welch periodogram method, which applied the Fast Fourier Transform (FFT) to windowed and segmented data to ensure smooth transitions. Additionally, we calculated the LF- HRV/HF-HRV ratio (LF/HF ratio) for exploratory analyses purpose.

Shapiro-Wilk tests were performed to assess the normality of the data. LF-HRV, HF-HRV and LF/HF ratio values were found to violate normality assumptions (*ps* < 0.001). Therefore, these values were log-transformed to normalize their distribution. Paired sample t-tests were conducted to compare participants’ CoDG and response times between the systolic and diastolic phases of the cardiac cycle. Pearson correlation analyses were performed to assess the relationships between HRV indices (RMSSD, LF-HRV, HF-HRV, LF/HF ratio), autistic traits, anxiety levels, and CoDG [systole-CoDG, diastole- CoDG, △CoDG (systole - diastole)]. Multiple regression analyses (MLR) were conducted to investigate the influence of HRV, autistic traits and anxiety levels on CoDG width. Participants were divided into high and low HRV groups based on the median-split of their RMSSD data. A two-way mixed-design analysis of covariance (ANCOVA) was calculated with cardiac cycle (systole vs. diastole) as a within- subjects factor and HRV group (High vs. Low) as a between-subjects factor, AQ and STAI scores were included as covariates to control for potential influences of autistic traits and anxiety levels on CoDG, basing on previous studies indicating that both autistic traits (measured by AQ) and anxiety traits (measured by STAI) can influence participants’ CoDG. Effect sizes were calculated using partial eta squared (η²). A *p*-value of < 0.05 was considered statistically significant.

## 3. Results

### 3.2 Behavioral results

A two-tailed *t*-test for the CoDG width indicated no significant differences between the systole (M = 7.59, SD = 1.96) and diastole (M = 7.51, SD = 1.77) conditions, *t*(34) = 0.49, *p* = 0.63, *d* = 0.08 (Fig. 3A). For the response times, the mean response time was 339.67 ms (SD = 66.26) in the systole trials and 327.15 ms (SD = 68.41) in the diastole trials, the mean reaction time was significantly longer in systole trials than in diastole trials, *t*(34) = 2.42, *p* < 0.01, *d* = 0.48 (Fig. 3B).

**Fig. 3.**
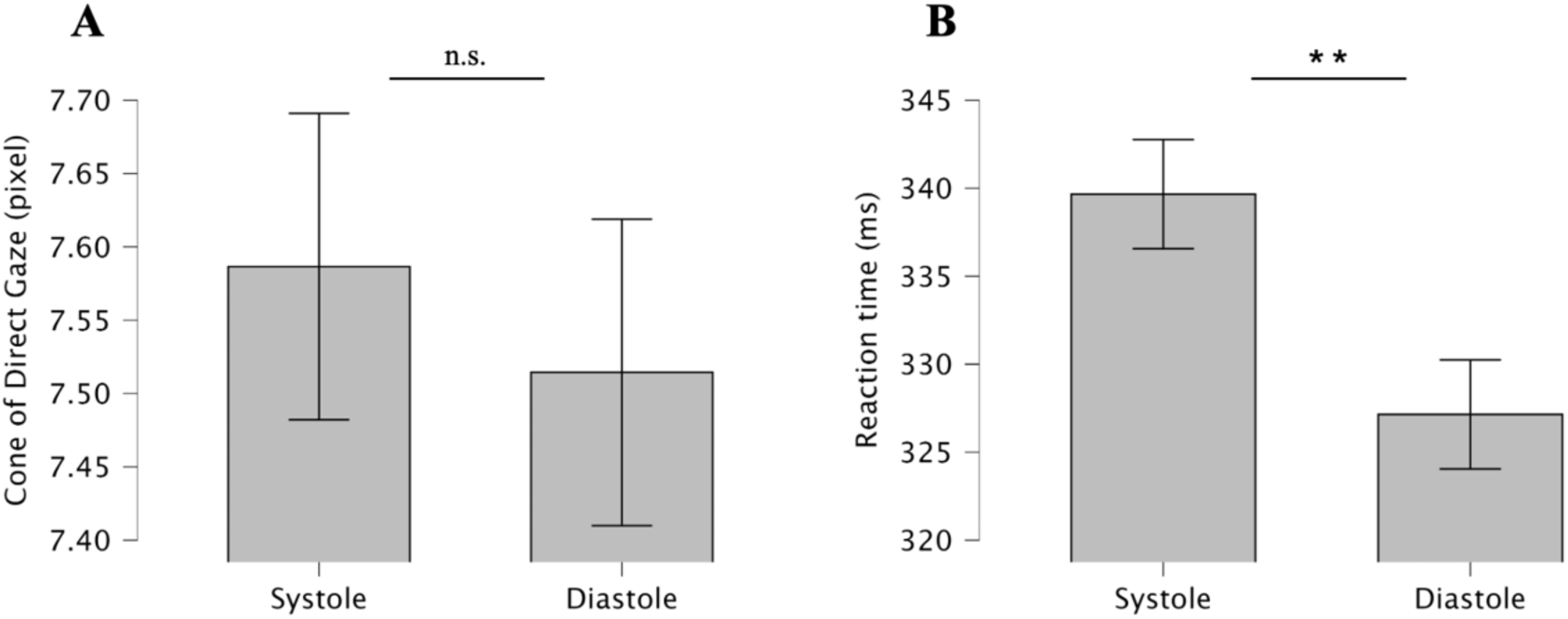
The mean cone of direct gaze width (**A**) and mean reaction time (**B**) for systole and diastole conditions. Error bars represent standard error of the mean (SEM). * * *p* < .01.

Pearson correlation analyses (Table 1) revealed that participants’ RMSSD were significantly correlated with both systole-CoDG (*r* = 0.43, *p* < 0.01) and △CoDG (systole - diastole) (*r* = 0.39, *p* = 0.020). Similarly, significant positive correlations were found between HF-HRV (log) and systole-CoDG (*r* = 0.44, *p* < 0.01), HF-HRV (log) and △CoDG (systole - diastole) (*r* = 0.34, *p* = 0.045) (see Fig. 4). No significant correlations were found between diastole-CoDG and RMSSD, LF-HRV (log), HF-HRV (log), or the LF/HF ratio (*p’s* > 0.06). No significant correlations were found between the AQ, STAI-s, or STAI- t scores and CoDG width [systole-CoDG, diastole-CoDG, or △CoDG (systole - diastole)] (*p’s* > 0.13).

**Fig. 4.**
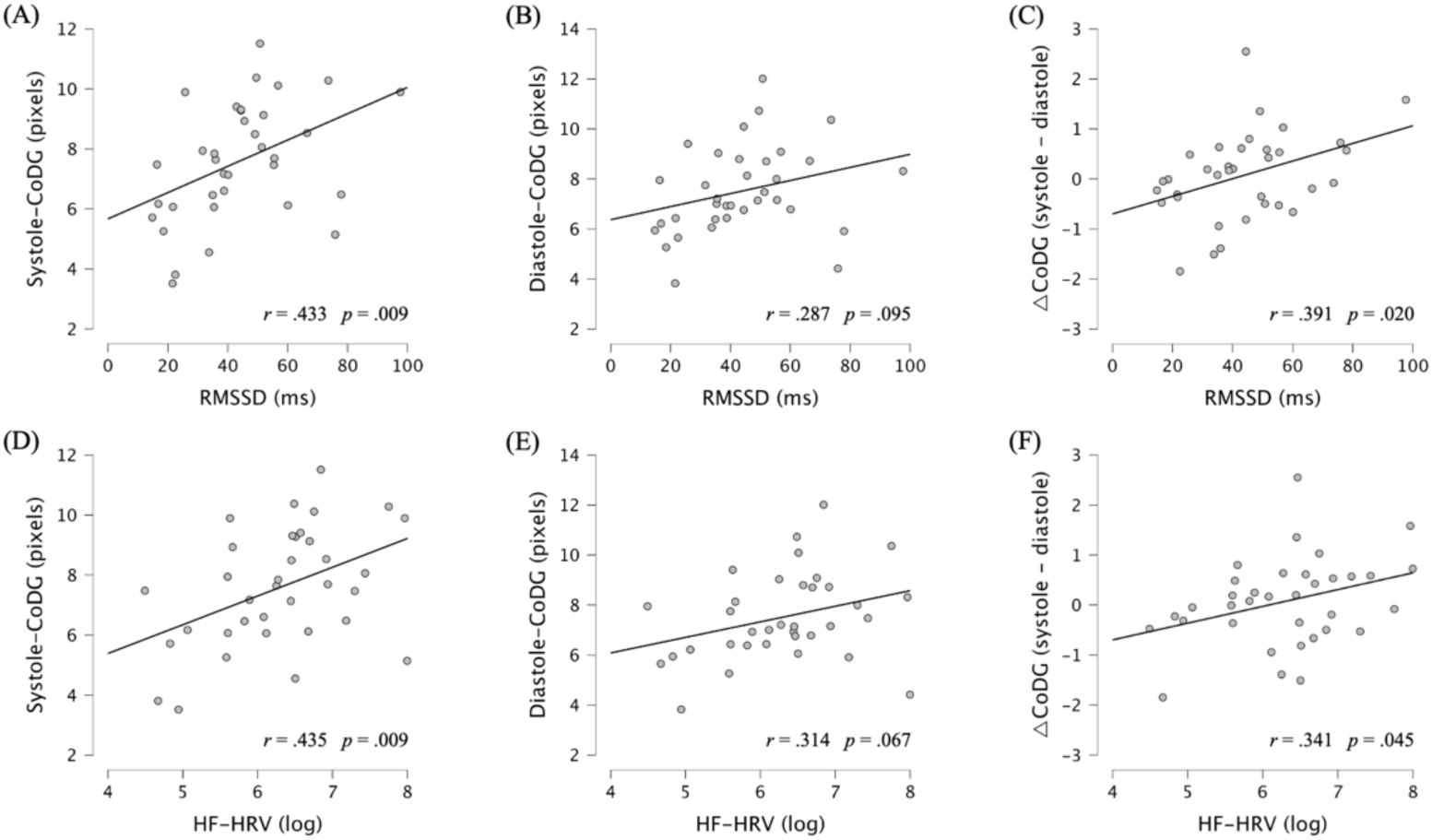
Correlations of root mean square of successive difference (RMSSD) and high-frequency heart rate variability (HF-HRV) with the width of the cone of direct gaze (CoDG) in systole condition (**A**, **D**), diastole condition (**B**, **E**) and the different width between systole and diastole conditions (**C**, **F**).

**Table 1:** Correlation coefficients between CoDG and HRV parameters.

|  | HRV parameters |  |  |  |
| --- | --- | --- | --- | --- |
|  | RMSSD | LF-HRV | HF-HRV | LF/HF Ratio |
| Systole-CoDG | 0.433 ** | 0.342 * | 0.435 ** | - 0.185 |
| Diastole-CoDG | 0.287 | 0.296 | 0.314 | - 0.093 |
| ΔCoDG (systole - diastole) | 0.391 * | 0.168 | 0.341 * | - 0.226 |

*Note.* RMSSD = root mean square of successive difference; LF-HRV = low-frequency heart rate variability; HF-HRV = high-frequency heart rate variability; LF/HF ratio = LF-HRV/HF-HRV ratio. \* $p < .05$ , \*\* $p < .01$ .

According to the results of the correlation analyses, a multiple linear regression was conducted to predict systole-CoDG and △CoDG (systole - diastole) from RMSSD, AQ, STAI-s and STAI-t scores. For systole-CoDG, the overall model was not significant [R² = 0.255, F(4, 30) = 2.57, *p* = 0.058], RMSSD was the only predictor that showed a significant positive association with systole-CoDG (β = 0.42, p = 0.014), while no significant effects were observed for AQ, STAI-s, or STAI-t scores (*p’s* > 0.17). For △CoDG (systole - diastole), the overall model was not significant [R² = 0.178, F(4, 30) = 1.63, *p* = 0.193]. RMSSD again emerged as a significant predictor (*β* = 0.43, *p* = 0.019), whereas AQ, STAI-s, and STAI- t did not predict △CoDG (*p’s* > 0.37).

Two-way mixed-design ANCOVA showed a significant interaction between cardiac cycle and HRV group, F(1, 30) = 6.90, *p* = 0.013, 𝜂^"^ = 0.187, and a significant main effect of HRV group, F (1, 30) = 10.37, *p* = 0.003, 𝜂^"^ = 0.257, on CoDG width. Additionally, neither main effects of cardiac cycle nor interaction effects between cardiac cycle and AQ, STAI-s, or STAI-t were significant (*p’s* > 0.52). For between-subjects effects, no significant main effects of AQ, STAI-s, or STAI-t in CoDG width were observed (*p’s* > 0.29). The results of the two-way mixed-design ANCOVA are shown in Table 2.

**Table 2.** Result of the two-way mixed-design ANCOVA for the effects of cardiac cycle and HRV group on CoDG width, controlling for AQ, STAI-s, and STAI-t.

| | df | F | p | $\eta_p^2$ |
| --- | --- | --- | --- | --- |
| Within-Subjects Effects |  |  |  |  |
| Cardiac cycle | 1 | 0.230 | 0.635 | 0.008 |
| Cardiac cycle × AQ | 1 | 0.415 | 0.524 | 0.014 |
| Cardiac cycle × STAI-s | 1 | 0.112 | 0.741 | 0.004 |
| Cardiac cycle × STAI-t | 1 | 0.384 | 0.540 | 0.013 |
| Cardiac cycle × HRV group | 1 | 6.902 | **0.013** | 0.187 |
| Between-Subjects Effects |  |  |  |  |
| AQ | 1 | 0.123 | 0.729 | 0.004 |
| STAI-s | 1 | 1.140 | 0.294 | 0.037 |
| STAI-t | 1 | 0.186 | 0.670 | 0.006 |
| HRV group | 1 | 10.373 | **0.003** | 0.257 |

*Note.* N = 35; HRV = heart rate variability; AQ = Autism-Spectrum Quotient; STAI-s = the state scale of the State-Trait Anxiety Inventory; STAI-t = the trait scale of the State-Trait Anxiety Inventory.

## 4. Discussion

In the present study, we used a modified gaze discrimination task to elucidate the effect of cardiac afferent signals on gaze perception. By presenting face stimuli with different gaze directions that coincided with either the systole or diastole phases of participants’ cardiac cycle, we calculated CoDG width for each cardiac phase separately. Contrary to our hypothesis, we did not observe direct cardiac cycle effects on the CoDG. Additionally, reaction times were significantly longer during cardiac systole than diastole. Moreover, we observed an interaction between cardiac cycle and HRV on CoDG. Specifically, the results showed that the higher the RMSSD, the wider the CoDG in the systole compared to the diastole condition, indicating that individuals with higher resting HRV have a better cardiac vagal control, which contributes to a heightened physiological context of self-reference during systole when making gaze direction judgments, leading to a wider range of gaze directions perceived as directed at them.

The result of paired t-test on cardiac cycle showed that there is no statistically significant difference in CoDG width in two cardiac conditions, suggesting that the manipulation of tasks did not consistently elicit a broadly effective cardiac cycle modulation. One likely explanation is that the short presentation time of the stimuli may be insufficient to trigger self-referential processes during the systole in all participants. Previous studies showed that CoDG was expanded by eliciting self-referential processes through external cues used longer stimulus presentation times compared to the present study, such as 600 ms (Stoyanova et al., 2010; Vida & Maurer, 2013) and 700 ms (Chen & Hietanen, 2024). In our study, the 100 ms stimuli duration was constrained by the transient nature of the cardiac cycle, in order to ensure that stimuli presentations were accurately timed to the specific cardiac phase and avoid potential floor or ceiling effects due to the presentation times. Although a duration of 100 ms is sufficient for processing gaze direction information (Bailey et al., 2014; Sato et al., 2007; Sato et al., 2016) and ensures timing with each phase of the cardiac cycle (Garfinkel et al., 2014), we could not entirely eliminate the impact of individual differences, such as interbeat interval length and the time required to trigger effective self- referential processes.

The results of correlation, regression, and ANCOVA analyses offered a more comprehensive perspective on the cardiac cycle effects. Specifically, higher vagally modulated HRV predicted a wider CoDG during systole, as well as the width of CoDG difference between systole and diastole conditions, while exhibiting no significant effect on CoDG width during diastole. Similarly, Izaki and colleagues reported a cardiac effect on social pain by time-locking the exclusion events to different cardiac cycle, finding that participants with higher RMSSD showed greater attenuation of pain in the systole condition relative to diastole, as well as a larger CoDG difference between the two cardiac conditions (Izaki et al., 2024). Additionally, another study indicated that response inhibition was better at systole, while participants with higher RMSSD tended to show reduced response inhibition during diastole (Rae et al., 2018). Systolic phase is the period during which arterial baroreceptors are activated, sensing and transmitting blood pressure and heartbeat information to the cortical level via the afferent pathway, which represents a peak in brain-heart communication during the fluctuations of cardiovascular activity. In parallel, the cardiac vagal tone, as assessed by HRV metrics such as RMSSD, reflects the trait-like cardioregulation through parasympathetic nervous system (Laborde et al., 2017). Higher HRV is associated with stronger baroreflex, which facilitates frequent heart rate adjustments through vagal efferent activity to maintain stable blood pressure (Pham et al., 2021). Taken together, individuals with higher HRV are likely to exhibit a more frequent and sensitive response to afferent signals from the heart and blood vessels, which may consequently contribute to a better cardiac modulation in social cognition.

A growing number of psychophysiological research have investigated the role of cardiac vagal tone in cognitive functions (for reviews, see Arakaki et al., 2023; Forte et al., 2019), such as studies have found HRV to be positively associated with decision-making (Forte et al., 2021; Forte et al., 2022; Prell et al., 2024), working memory (Hansen et al., 2003; Zeng et al., 2023), emotion regulation (Balzarotti et al., 2017; Mather & Thayer, 2018; Williams et al., 2015) and compassion (Di Bello et al., 2020). Notably, emerging evidence also demonstrates a strong link between HRV and interoceptive accuracy (Blickle et al., 2024; Lischke et al., 2021; Rominger et al., 2021; Vabba et al., 2023), which refer to the objective ability to detect one’s own internal bodily signals (Garfinkel et al., 2015). Individuals with high interoceptive accuracy are more sensitive to their internal bodily cues, leading to more intense emotional experiences (Parrinello et al., 2022) and heightened arousal (Dunn et al., 2010; Pollatos et al., 2007). von Mohr et al. (2021) measured participants’ interoceptive accuracy, rather than HRV, and showed that individuals with high interoceptive accuracy exhibited greater emotional egocentricity during systole, suggesting cardiac afferent signals accentuated self-referential processing when receiving information about the other’s emotional state. In the work by Ambrosini et al. (2019), researchers employed ambiguous morphed images combining self and other faces and revealed that stimuli presented during systole significantly accelerated reaction times for identifying as ‘self-faces’ compared to diastole.

Likewise, the present study used rapidly presented gaze stimuli from various directions to create ambiguous judgment conditions at specific angles (e.g., 3 pixels left, direct gaze, and 3 pixels right), aiming to investigate the reference cues participants rely on for gaze direction judgment across different phases of baroreceptor activity. Participants with higher HRV may implicitly use the cardiac afferent signals as a self-referential anchor in social decision-making during systole, leading to a broader range of self-directed gaze perception.

Our study also found that HF-HRV exhibited correlation patterns with CoDG similar to those observed between RMSSD and CoDG. Specifically, HF-HRV was positively correlated with CoDG during systole and with the CoDG difference across cardiac phases. Both HF-HRV and RMSSD serve as indexes of parasympathetic modulation of heart rate (Malik et al., 1996; Thomas et al., 2019). However, unlike RMSSD, HF-HRV is not considered a pure vagal tone indicator, since it reflects heart rate changes related to the respiratory cycle (Grossman & Taylor, 2007). As such, HF-HRV is also used to capture respiratory sinus arrhythmia (RSA) and is called the ‘respiratory band’ (Javorka et al., 2020; Laborde et al., 2017; Shaffer & Ginsberg, 2017). Given that participants’ respiratory rates were neither measured nor controlled in this experiment, evaluating respiration-adjusted HF power could provide a more informative perspective for studies examining the impact of autonomic reactivity on cognitive functions. Interestingly, we found a significant correlation between LF-HRV and CoDG during systole. LF-HRV has traditionally been considered an index of cardiac sympathetic activity, however, these perspectives have faced increasing challenges as growing evidence suggests that LF-HRV may reflect baroreflex function rather than cardiac sympathetic tone (Goldstein et al., 2011; Moak et al., 2007; Rahman et al., 2011; Reyes del Paso et al., 2013; but see Martelli et al., 2014). These contradictory findings underscore the necessity for more refined control, including manipulations (Bernardi et al., 2000; Duschek et al., 2009; Mukai & Hayano, 1995) and assessments (Clark et al., 2018; Li & Zheng, 2022) of participants’ sympathetic activation, in future studies.

Despite the null result for the cardiac cycle on CoDG, paired t-test results demonstrated that reaction times were significantly longer for stimuli presented during systole compared to those presented during diastole, consistent with the findings of previous studies (McIntyre et al., 2008; Ren et al., 2022; Saltafossi et al., 2023; Yang et al., 2017). The systolic inhibition hypothesis suggests that systolic baroreceptor activity induces neural noise, which interferes with the processing of non-affective exteroceptive stimuli, including visual, auditory, and somatosensory domains (Al et al., 2020; Edwards et al., 2002; McIntyre et al., 2006; Walker & Sandman, 1982; Wilkinson et al., 2013; Schulz et al., 2009; Schulz et al., 2020; for a review, see Skora et al., 2022). In line with this notion, gaze stimuli repeatedly delivered in phase with cardiac afferent signals in the present study were less effectively detected than those in diastolic phases, which may in turn increase the reliance on internal cues, as interoceptive signals are accentuated during systole (Critchley & Harrison, 2013; Nokia et al., 2024).

We ruled out the possible effects of individual traits (anxiety levels and autistic traits) in the present study. Our findings indicated no significant association between participants’ STAI scores and CoDG to neural facial stimuli, which is consistent with observations of Hu et al. (2017), but contrary to some prior studies (Chen et al., 2017; Harbort et al., 2013; Harbort et al., 2017; Schulze et al., 2013) that demonstrated higher anxiety levels were associated with widened CoDG in response to non-emotional facial expressions. It is noteworthy that these studies used clinical samples with social anxiety disorder (Harbort et al., 2013; Harbort et al., 2017) or different questionnaires (Chen et al., 2017; Schulze et al., 2013) such as the Social Phobia Scale (SPS, Mattick and Clarke, 1998) and the Social Phobia Inventory (SPIN; Connor et al., 2000), which differ from those used in the present study. We also found no effect of autistic traits on CoDG, which is consistent with Williams et al. (2023) but contrary to Matsuyoshi et al. (2014), which found that higher autistic traits were associated with narrower CoDG. To date, studies examining the effects of autistic traits on CoDG are scarce and their findings are contradictory. Moreover, based on the AQ cut-off value of 33 in the Japanese sample (Wakabayashi et al., 2006), only two participants in this study reached the high autistic traits threshold. Therefore, the findings from this study may not accurately reflect the effect of autistic traits on CoDG. Furthermore, as previously discussed, it is possible that the unique design of the paradigm contributed to the absence of a significant effect of individual traits on CoDG with neutral facial stimuli. Nonetheless, these findings emphasize the robustness and stability of cardiac effects on gaze perception across individual differences.

## 5. Conclusion

In conclusion, the present study used a modified gaze discrimination task and found that cardiac afferent signals and cardiac vagal tone affect gaze perception. Specifically, individuals with higher RMSSD showed a wider CoDG in the systole condition than in the diastole condition. Moreover, this effect was not influenced by individual traits such as anxiety levels or autistic traits. This study provides the first empirical evidence for the influence of cardiac cycle and parasympathetic activity on gaze perception, highlighting the role of interoceptive signals in social judgment.

